# Recalibrating differential gene expression by genetic dosage variance prioritizes functionally relevant genes

**DOI:** 10.1101/2024.04.10.588830

**Authors:** Philipp Rentzsch, Aaron Kollotzek, Pejman Mohammadi, Tuuli Lappalainen

**Author notes:** Max Delbrück Centre for Molecular Medicine, Berlin, Germany (Aaron Kollotzek).

## Abstract

Differential expression (DE) analysis is a widely used method for identifying genes that are functionally relevant for an observed phenotype or biological response. However, typical DE analysis includes selection of genes based on a threshold of fold change in expression under the implicit assumption that all genes are equally sensitive to dosage changes of their transcripts. This tends to favor highly variable genes over more constrained genes where even small changes in expression may be biologically relevant. To address this limitation, we have developed a method to recalibrate each gene’s differential expression fold change based on genetic expression variance observed in the human population. The newly established metric ranks statistically differentially expressed genes not by nominal change of expression, but by relative change in comparison to natural dosage variation for each gene. We apply our method to RNA sequencing datasets from rare disease and in-vitro stimulus response experiments. Compared to the standard approach, our method adjusts the bias in discovery towards highly variable genes, and enriches for pathways and biological processes related to metabolic and regulatory activity, indicating a prioritization of functionally relevant driver genes. With that, our method provides a novel view on DE and contributes towards bridging the existing gap between statistical and biological significance. We believe that this approach will simplify the identification of disease causing genes and enhance the discovery of therapeutic targets.

## INTRODUCTION

Since the advent of cDNA microarrays (Schena et al. 1995), differential gene expression profiling has been used to examine the characteristic gene regulatory changes of a specific phenotype, disease state or perturbation. Statistical tests are used to determine whether the observed differences in expression of each single gene between groups of samples are statistically significantly differentially expressed (DE) over a chosen statistical threshold. Over the years, many different bioinformatics methods to test DE have been developed, adapting rigorous statistical tests to the particularities and assumptions of DNA microarrays and later RNA-Seq. Some of the currently most popular methods for differential gene expression analysis are DESeq2 (Love et al. 2014), edgeR (Robinson et al. 2010), and limma-voom (Ritchie et al. 2015). These tools are designed to work with the count-based nature of RNA-Seq data and incorporate normalization procedures to account for differences in sequencing depth and RNA composition across samples. Like most statistical tests, DE methods have been shown to sometimes produce false positives, particularly when analyzing RNA-Seq datasets of large sample sizes. To address this issue, several different approaches of multiple testing correction have been developed (Benjamini and Hochberg 1995; Ge et al. 2009; Ignatiadis et al. 2016) and are applied in many of the popular methods.

While DE testing quantifies the statistical significance of differential expression, it is agnostic to its *biological relevance*, i.e. whether the detected change in gene expression meaningfully reflects or contributes to changes in cellular functions. This means that a test’s p-value does not carry any inherent meaning (Wasserstein and Lazar 2016; Greenland et al. 2016), and well-powered DE studies can result in hundreds if not thousands of differentially expressed genes. As a consequence, the actual expression fold change - the ratio of the expression levels between the two sample groups - is often considered an important secondary parameter (Cui and Churchill 2003; McCarthy and Smyth 2009; Zhang and Cao 2009; Jung et al. 2011; Harrison et al. 2019). Many DE studies apply an arbitrarily chosen minimum fold change threshold and inspect only statistically significant genes surpassing this cut-off. This makes sense insofar that the magnitude of an expression change is obviously relevant (Naqvi et al. 2023) and for a single gene bigger expression changes tend to matter more than smaller ones. However, the absolute fold change in itself is of limited relevance when comparing changes between different genes, because different genes have different levels of dosage constraint (Rice and McLysaght 2017). This means that a measured change in expression in one gene may be comparable to biological noise and may fall within the spectrum of natural variation in the population. However, a deviation of the same magnitude in another gene can be highly unusual and lead to immediate cellular consequences, either by affecting other genes or by directly altering the phenotype. Such a change could hence be described as not only statistically but also biologically significant. Prior work shows that genes with constrained expression are enriched for drivers of cellular processes and disease (Lek et al. 2016; Rice and McLysaght 2017; Mohammadi et al. 2019; Karczewski et al. 2020; Collins et al. 2022; Dong et al. 2023). It follows that the subsequent difference in gene responsiveness makes nominal fold change a poor proxy for biological relevance, an effect that may contribute to the systematic differences between gene expression changes and associations between genetic variants and traits (Mostafavi et al. 2023).

The best studied quantity that describes how tolerant genes are to dosage variation is haploinsufficiency, the gene’s intolerance to heterozygous deletion or loss-of-function variants. A related concept of triplosensitivity refers to intolerance to duplication. Beyond these large changes that affect one entire copy of a gene, especially non-coding genetic variants affect the expression of nearby genes (Flynn and Lappalainen 2022). For many genes, as recently exemplified for the transcription factor SOX9 (Naqvi et al. 2023), these regulatory changes are associated with differences in phenotype and disease risk (Albert and Kruglyak 2015). To our knowledge, there exist currently no assays that measure the viable dosage range of mRNA expression for each gene in a genome. However, it has been shown that haploinsufficiency of genes aligns well with the amount of purifying selection in the genome (Lek et al. 2016; Karczewski et al. 2020; Collins et al. 2022). Indeed, depletion of genetic variation in populations has been a powerful genome-wide approach for detecting molecular variation that is not tolerated by natural selection.

Here, we established a novel calibration method to measure differential expression by scaling standard DE testing with the population genetic dosage variation estimate V^G^ that we have introduced earlier. We tested this approach in multiple previously published datasets, and showed that the resulting recalibrated gene expression fold changes removed the DE discovery bias towards highly variable genes, and that selecting DE genes by recalibrated fold change increases the enrichment of regulatory and metabolic GO terms, i.e. potential driver genes of cellular processes. In order to apply recalibration to as many genes as possible, we trained a machine learning model based on other genetic constraint metrics. This enabled us to expand recalibration analysis not only to more genes, but also perform tissue-specific recalibration. Analysis of brain expression data in autism spectrum disease indicated increased enrichment of relevant cellular processes and highlighted previously known disease genes. In summary, our study demonstrates that measuring differential gene expression in nominal expression fold change is affected by varying levels of dosage tolerance and may overlook relevant driver genes. The fold change recalibration method established here resolves this issue by measuring expression changes in terms of deviation relative to the natural variability range. Our observations suggest that this recalibration approach improves the focus of DE testing and has the potential to enrich DE results for biologically significant genes.

## RESULTS

### Variance in gene expression as a fold change recalibration metric

The amount of population variability in expression is different for each gene. While some of this variability is due to technical factors, physiological differences, or external influences from the environment, some of it is due to genetic differences between individuals. To infer the component of natural variation that is due to genetic *cis*-regulatory differences, we previously introduced ANEVA, a method for assessing the gene expression distribution based on allelic expression (AE, Mohammadi et al. 2019). Applying ANEVA to GTEx v8, an RNA-Seq dataset from more than 800 individuals, we derived an estimate of gene expression variance (V^G^) for each human gene. Since both expression and variability depend on body tissue and cell type, V^G^ is initially calculated separately for 49 different tissues in GTEx. An overall V^G^ value is then calculated as the weighted harmonic mean across tissues.

As V^G^ is based on the allelic fold change (Mohammadi et al. 2017) of genetic regulatory effects, its unit is analogous to log fold changes used in DE analyses. The magnitude of V^G^ describes the range in which observed gene expression changes fall within the population (Fig. 1A). Large values imply greater variability and thus a wider range of expression levels in the population. V^G^ hence serves as a parameter to classify an expression change relative to the respective gene’s mean, i.e where a sample with the observed change would fall within the natural (population) range.

**Figure 1:**
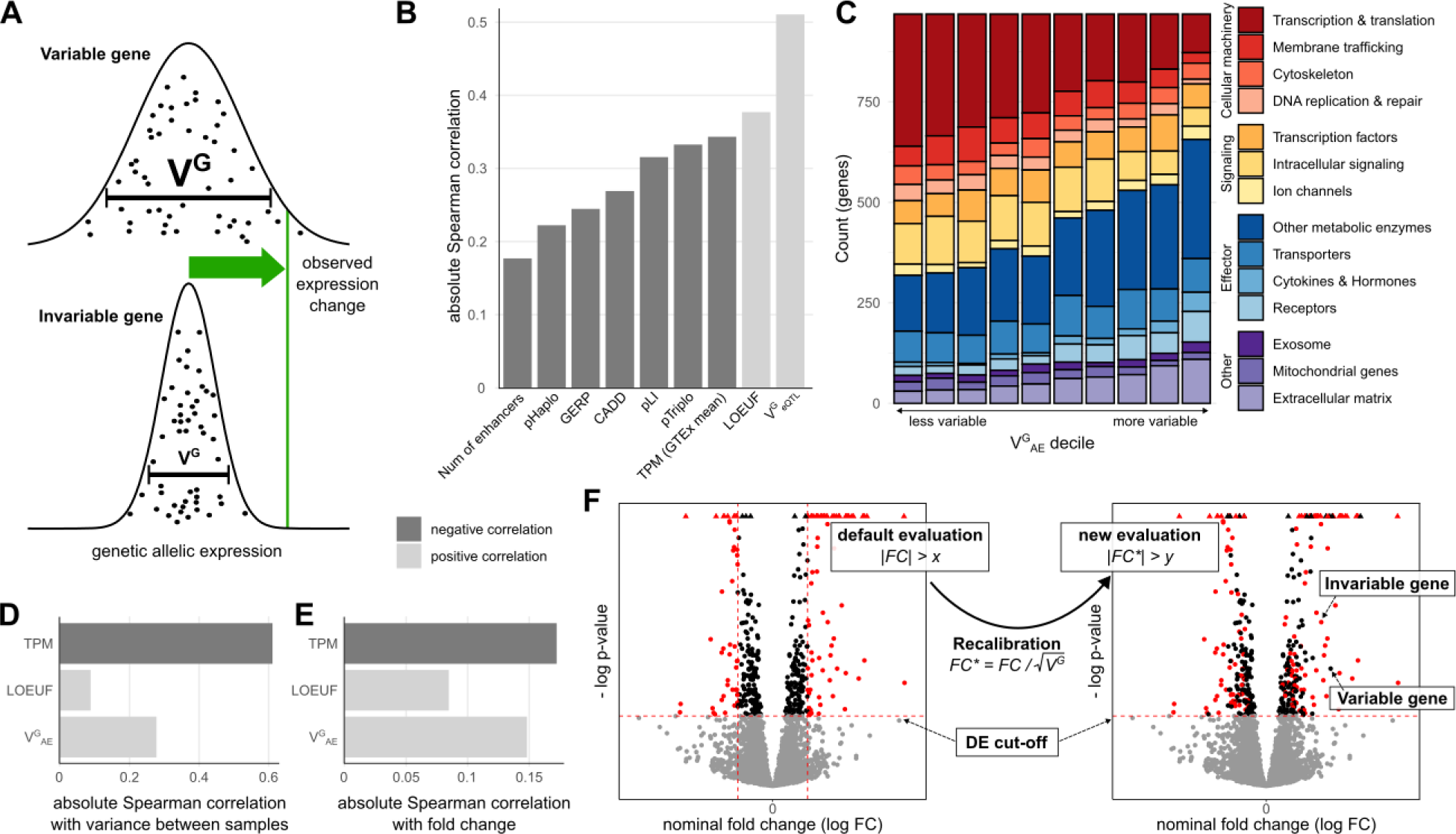
Fold change recalibration using V^G^. **A)** Depending on variability of a gene, an observed fold change may be within the natural variation that is observed in the general population. V^G^ is an estimate of this population variance. **B)** Correlation of V^G^ with other gene metrics. Negative correlations are colored dark gray, while positive correlations are light gray. **C)** Distribution of functional categories in genes split by V^G^ decile. Some functional categories like transcription and translation associated genes were enriched for small V^G^ estimates while others like receptors were enriched for high V^G^ estimates. **D)** Correlation of gene metrics with experimental gene expression variance. Data sourced from Alasoo et al. studying the effects of IFNγ treatment on 81 human macrophage samples. **E)** Correlation of gene metrics with absolute gene expression changes in the IFNγ experiment. **F)** Concept of gene expression recalibration using V^G^. As a result of this process, genes are prioritized not by nominal log fold change (FC) but by recalibrated log fold change (FC*).

In our previous publication *Mohammadi et al.* 2019, we showed that the V^G^ of a gene is correlated with selective constraint. Known haploinsufficient genes have lower V^G^ on average and V^G^ is correlated with other constraint metrics such as conservation scores or the probability of being loss-of-function intolerant (pLI, Fig 1B). Here, we measured its correlation to additional gene constraint metrics. One of the most highly correlated metrics (⍴_Spearman_=0.38) is the loss of function tolerance metric LOUEF (Karczewski et al. 2020). Most interesting are the haploinsufficiency and triplosensitivity metrics pHaplo and pTriplo (Collins et al. 2022). While V^G^ is less correlated with both metrics than, for example, LOEUF, we found that, in contrast to the latter, V^G^ is more correlated with pTriplo (⍴_Spearman_=-0.33) than pHaplo (⍴_Spearman_=-0.22), indicating its potential in capturing sensitivity to both up- and downregulation.

In addition to metrics of constraint, we found that V^G^ estimates are associated with gene function. As described for a simpler GTEx-derived ‘allelic Fold Change’ metric (Dong et al. 2023), genes annotated as part of the central cellular processes, particularly those involved in transcription and translation, are enriched among genes with low V^G^. In contrast, effector proteins such as receptors, as well as extracellular matrix proteins, are more likely to have a high V^G^ (Fig. 1C).

We next investigated how V^G^ is correlated to gene expression variance in experimental data. To this end, we analyzed a dataset of 81 human macrophage samples that were treated with IFNγ (Alasoo et al. 2018). As expected due to the well-documented mean-variance relationship, expression level (transcripts per million, TPM) was highly correlated with inter-individual variance between control samples, but also V^G^ and LOEUF were correlated to variance (Fig. 1D), which is not unexpected for different metrics of population variance. More notably, the expression fold change upon IFNγ stimulus was also correlated with V^G^ (Fig. 1E, ⍴_Spearman_=0.148, 95%-confidence interval (CI_95_)=[0.130;0.166]), significantly more so than LOEUF (⍴_Spearman_=0.085, CI_95_=[0.066;0.103]) and within the range of TPM (⍴_Spearman_=-0.172, CI_95_=[-0.190;-0.155]). While DE significance testing accounts for the variance in the studied dataset, the correlation between fold change and V^G^ indicates that genes that are more likely to be noticeable outliers in a DE study also have a higher population variance in general.

The idea behind the original ANEVA method was to have a test that evaluates whether the genetic effect in a gene in one particular sample is an outlier compared to the genetic variance in the population. Conversely, here we propose to use the derived variance metric V^G^ to rescale non-genetic gene expression differences between two groups in order to compare expression changes across different genes on the same scale, relative to the population variance in each. While the above findings provide support for using V^G^ to rescale gene expression fold changes, the main motivation for V^G^ as a metric itself is unit equivalence. Unlike scores such as LOEUF, GERP, CADD or pLI, where there is no direct relationship between score value and variability, a gene with a V^G^ of 0.02 is twice as variable as a gene with a V^G^ of 0.01. We used this property to recalibrate the experimental log fold change, by scaling it by the average change in expression. This average change in expression is the standard deviation, which is the square root of the variance. Thus, the approach, which we call recalibration, we standardize the observed fold change for each gene with the standard deviation of the genetically regulated gene expression given by the square root of V^G^ (Fig. 1F). To distinguish between pre- and post-recalibration, we refer to log fold changes obtained from an experiment as ‘nominal’ and fold changes after rescaling as ‘recalibrated’. Recalibration is an additional analysis step after significance testing, and it changes the relative order of genes assigned as significant based on standard DE testing. This means that small changes in expression for some genes may be considered relatively more significant than the same change would be for other genes.

### Recalibration shifts the analysis focus

To assess the effect of a recalibration with V^G^, we used the previously introduced dataset of 81 human macrophage samples stimulated with IFNγ. After processing the samples and selecting DE genes with a strict false discovery rate of less than 0.001, the dataset yields 20,402 DE genes (out of 29,438 in the dataset). Of these, 5,573 genes also have an absolute FC greater than 1. The dataset hence serves as an excellent illustration of the need for selection criteria beyond statistical significance to identify the most relevant genes for further analysis. To compute recalibrated fold changes, we restricted further analysis to the 16,282 genes in the dataset for which V^G^ estimates have been calculated. Of these, 13,372 were DE and 3,317 DE genes had an aFC greater than 1. The recalibrated fold changes were highly correlated with the nominal fold changes (Pearson correlation of absolute values= 0.84, Fig. 2A).

**Figure 2:**
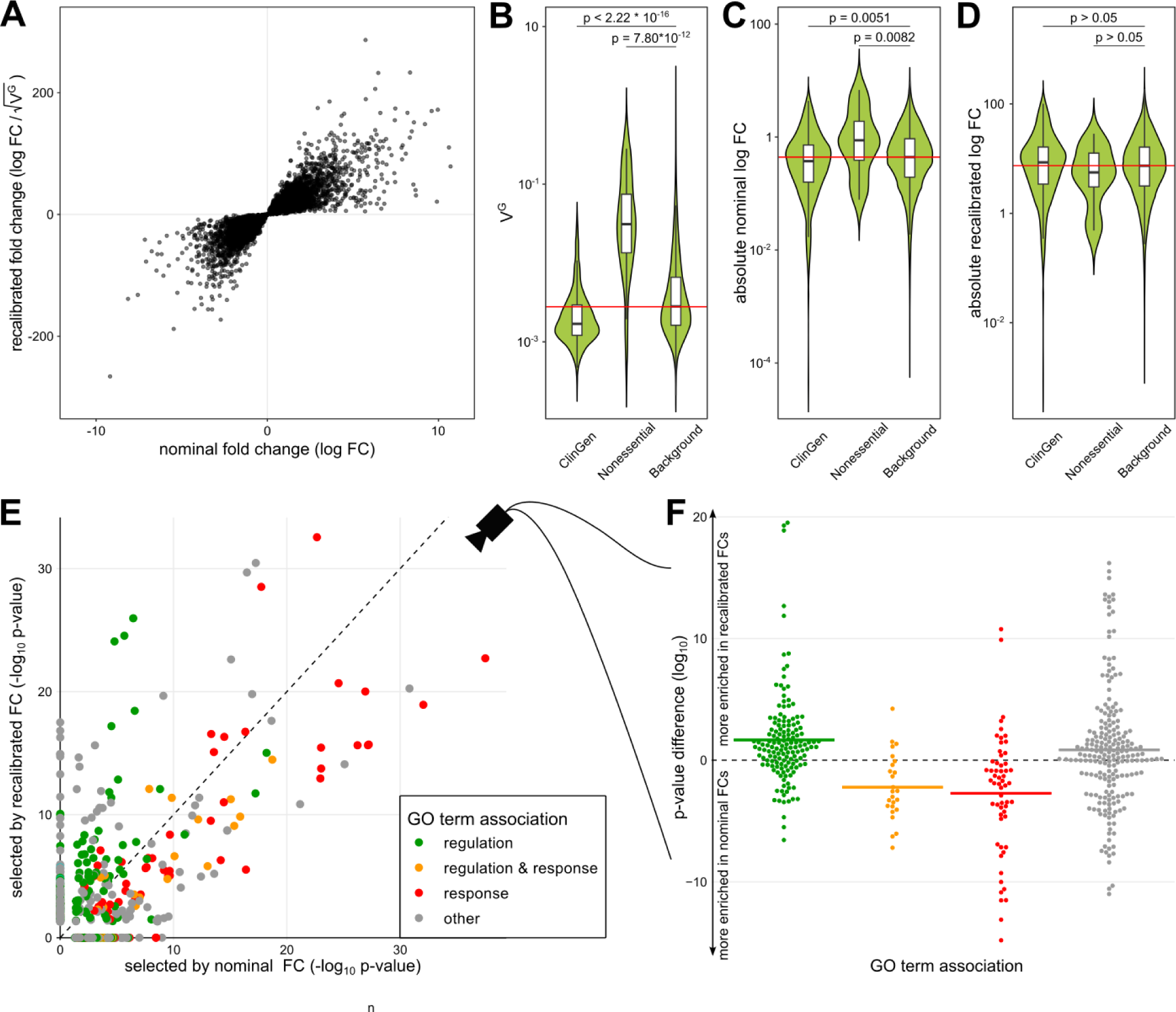
Fold change recalibration of a DE experiment of IFNγ treatment. **A)** Nominal compared to recalibrated fold changes of individual genes from *Alasoo et al. 2018*. **B-D)** Impact of recalibration on genes from three different gene sets: ClinGen, Nonessential and all other genes (Background). **B)** displays the distribution of V^G^ per set, **C)** absolute log fold changes before recalibration (nominal) and **D)** the recalibrated absolute log fold changes. Each violin represents the total distribution of values, with boxes in the violin indicating 25% and 75% quantiles and the median illustrated as a black line. Whiskers are 1.5 times the interquartile range. The red line denotes the median over all genes. Significance labels are the result of a two-sided comparison with Mann-Whitney U test of the two groups. **E)** Effect on GO term enrichment: Adjusted p-values of GO term enrichment from 2,000 genes selected by nominal compared to recalibrated FC. GO-terms are colored based on string matching (association). Center line marks the same enrichment in both sets. GO terms above the line are more enriched before the recalibration while terms below the line are more enriched after. **F)** Different view on the change in GO term enrichment p-values grouped by associations. Colored dots are GO terms while the colored vertical lines are the log-mean per association. Terms containing the word ‘regulation’ (green) are on average more enriched after recalibration, while terms containing the word ‘response’ (red) are on average more enriched before recalibration.

To examine the effect of the recalibration, we first turned to sets of well-characterized genes. Known haploinsufficient genes from ClinGen on median had lower than average V^G^ estimates, while those of non-essential genes were generally higher than average (Fig. 2B). Non-essential genes had higher absolute fold changes upon IFNγ stimulus than ClinGen and background genes, indicating that a standard DE analysis might easily focus on these functionally irrelevant genes (Fig. 2C). The recalibration results in an adjustment of the fold change ranges for the different gene sets (Fig. 2D), with their respective distributions no longer being significantly different.

Using the 2,000 most differentially expressed genes by nominal and recalibrated absolute FC, respectively, we observed a shift in GO term enrichments (Fig. 2E, Supplemental Table S1). In particular, terms matching the string ‘response’, like ‘immune response’ (GO:0006955) and ‘response to other organism’ (GO:0051707), were less enriched, while terms matching ‘regulation’, like ‘regulation of biological process’ (GO:0050789), were more enriched among the genes selected after recalibration (Fig. 2F). This effect was robust to the number of genes analyzed (Supplemental Fig. S1) or when the gene set enrichment was calculated based on gene ranks (GSEA, Supplemental Fig. S2). GO term enrichments are not independent of each other as most genes are associated with many different GO terms. Clustering the GO terms by the genes associated with each term, we found that the two clusters enriched for GO terms whose names contain the word ‘response’ are on average less enriched after recalibration, while 6 out of 7 GO term clusters where multiple terms match the word ‘regulation’ are more enriched after recalibration (Supplemental Fig. S3). However, it should be noted that while the differences in enrichment can be large and only 1,470 genes are in the top 2,000 in both lists, the change in the number of genes per GO term is small (Supplemental Fig. S4). We found the same enrichment of GO terms matching ‘regulation’ and also of those matching the term ‘metabolic’ when performing the same analysis on experiments of macrophages perturbed with Salmonella and Salmonella+IFNγ from the same experimental data source (Supplemental Fig. S5). These results indicate that nominal fold change captures *response* to stimuli, while recalibration enriched for *regulators* of cellular response.

Next, we wanted to analyze whether this pattern holds for a DE study of a more limited size than the well-powered *Alasoo et al. 2018* data. In this notion, we analyzed the data from *Findley et al. 2021* with multiple in vitro stimuli of 3 different cell lines. Here, our focus fell on the response in expression of metallothionein genes due to a perturbation of cells with copper ion solution. Metallothioneins are a family of proteins that have the ability to bind metal ions, providing protection against metal toxicity (Sekovanić et al. 2020). In the experiment, a total of 9 metallothionein genes are significantly upregulated with fold changes between 2 and 7 (Supplemental Fig. S6A) and the associated GO term ‘cellular response of copper ion’ (GO:0071280) is significantly GO enriched. Recalibration reduces this enrichment (Supplemental Fig. S6B), as the VG values of the metallothionein genes are in the top quartile of all genes. We make the same observation for perturbation with zinc ion solution (Supplemental Fig. S7). Thus, the choice of DE approach depends on the goals of the analysis: the standard approach readily picks up genes that respond to the given stimulus. However, recalibration provides a gradual adjustment to DE gene rankings, which may help to deprioritize highly variable responder genes to highlight molecular drivers of cellular response.

### Expanding recalibration to more genes via other gene metrics

A challenge in the recalibration approach is that V^G^ estimates based on allelic expression data (V^G^_AE_) are currently missing for many genes due to sparsity of allelic expression data. While this problem will be alleviated with growing datasets, we have previously established an alternative method of calculating expression variance based on eQTLs (V^G^_AE_). These have been generated from GTEx data for 24,685 genes (compared to 16,282 genes with V^G^_AE_, Fig. 3A). These V^G^_AE_ estimates are highly correlated (⍴_Spearman_ = 0.48, Fig. 1B), and eQTL-based V^G^_AE_ for recalibration resulted in a similar trend of raising regulatory over response GO terms comparable, with generally higher enrichment p-values due to the larger number of genes (Fig. 3B, Supplemental Fig. S8). However, V^G^_eQTL_ is strongly correlated with the GTEx TPM (⍴_Spearman_ = -0.76, compared to ⍴_Spearman_ = -0.34 for V^G^_AE_). While some of this correlation is likely due to biology, with highly expressed genes being more constrained (Lek et al. 2016), the high correlation for V^G^_eQTL_ may derive e.g. from differences in the power of eQTL calling. This, and the fact that AE data is naturally unaffected by environmental and technical factors, motivated the primary use of V^G^_AE_ for recalibration.

**Figure 3:**
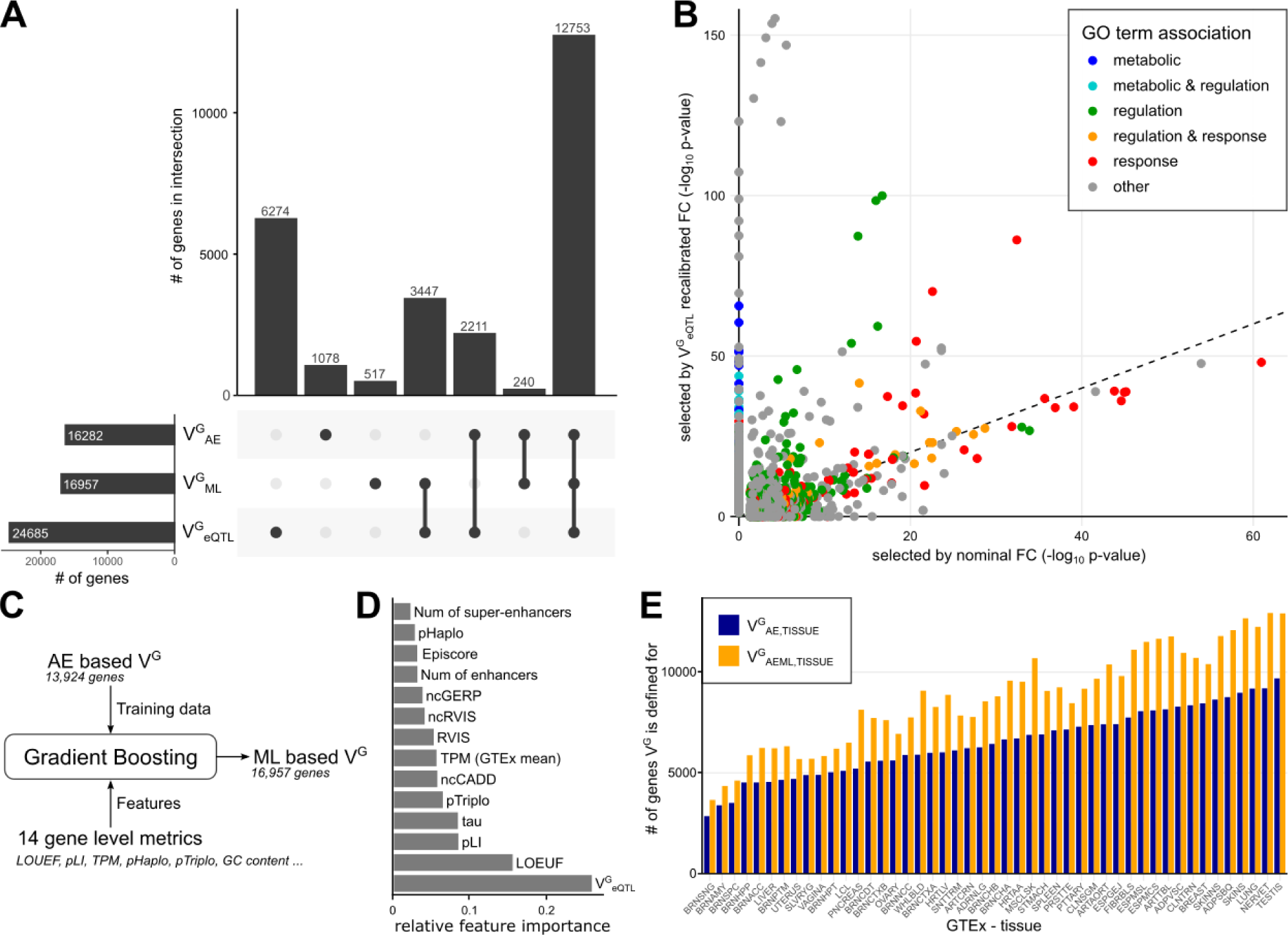
Recalibration with other metrics. **A)** The number of genes with different versions of V^G^ estimates. **B)** Impact of recalibration on GO term enrichment in the IFNγ stimulus dataset when using eQTL-derived V^G^. **C)** The concept of training a machine learning model of V^G^. **D)** Importance of different gene metrics for the machine learning model. **E)** Number of genes with V^G^ predicted per GTEx tissue, with ML and without.

To predict V^G^-like estimates for additional genes, we sought to leverage its high correlation with other metrics. We used a straightforward machine learning (ML) approach using gradient boosting trees to train predictive models using V^G^_AE_ as a label (Fig. 3C). K-nearest neighbor feature imputation was used to maximize the number of genes to be predicted, ultimately enabling the prediction of 16,957 genes (Fig. 3A). The best model achieved a log root mean square error of 0.77 on the hold-out test set and a Spearman correlation of 0.64 with V^G^_AE_ (Supplemental Fig. S9), using V^G^_eQTL_ and LOEUF as most important model features (Fig. 3D, Supplemental Table S2). The metric derived from the model, V^G^_ML_, has the same unit as V^G^_AE_ and can be used interchangeably for recalibration. V^G^_ML_ covered a smaller absolute range and had a lower variance between genes than V^G^_AE_ (variance of log values: 0.079 compared to 0.200 for V^G^_AE_). This resulted in a higher correlation between nominal and recalibrated FCs (e.g., ⍴_Pearson_=0.95 compared to 0.84 in Fig. 2A). While it follows that the changes in the GO term enrichments are smaller as well, the same general recalibration effects as for V^G^_AE_ were observed (Supplemental Fig. S10). Finally, we joined V^G^_AE_ and V^G^_ML_ scores in a single metric, V^G^_AEML_, that uses ML derived scores where V^G^_AE_ is missing in order to maximize the number of genes in recalibration, which ultimately covered a total of 20,246 genes.

In addition to the mean V^G^ values across tissues that we have used thus far, we used the same machine learning strategy to learn tissue-specific models for V^G^_AE_ of 47 GTEx tissues, using TPM and V^G^_eQTL_ as tissue-specific features. The trained models added V^G^ values for an average of 2,275 genes (range: 807 to 3,807, Fig. 3E, Supplemental Table S3), a mean increase of 34% (range: 16% to 56%). To cover as many genes as possible, we combined V^G^_AE,TISSUE_ and V^G^_ML,TISSUE_ into V^G^_AEML,TISSUE_.

### Tissue-specific expression fold change recalibration

Tissue-specific V^G^ estimates allow for experiment-specific recalibration. To explore this, we used the psychENCODE dataset (Gandal et al. 2018), which studied autism spectrum disorder (ASD), schizophrenia (SCZ) and bipolar disorder (BD) across hundreds of RNA-Seq samples primarily from the prefrontal cortex. The potential benefit of recalibration with tissue-specific rather than cross-tissue population variance is exemplified by significantly (binomial test p-value of 5.2 * 10^-4^) increased fraction of previously known ASD genes (An et al. 2018) having their lowest V^G^ in one of the 14 GTEx brain tissues (Fig. 4A). This indicates that V^G^, unlike genetics-based scores, can capture tissue-specific regulatory constraint.

**Figure 4:**
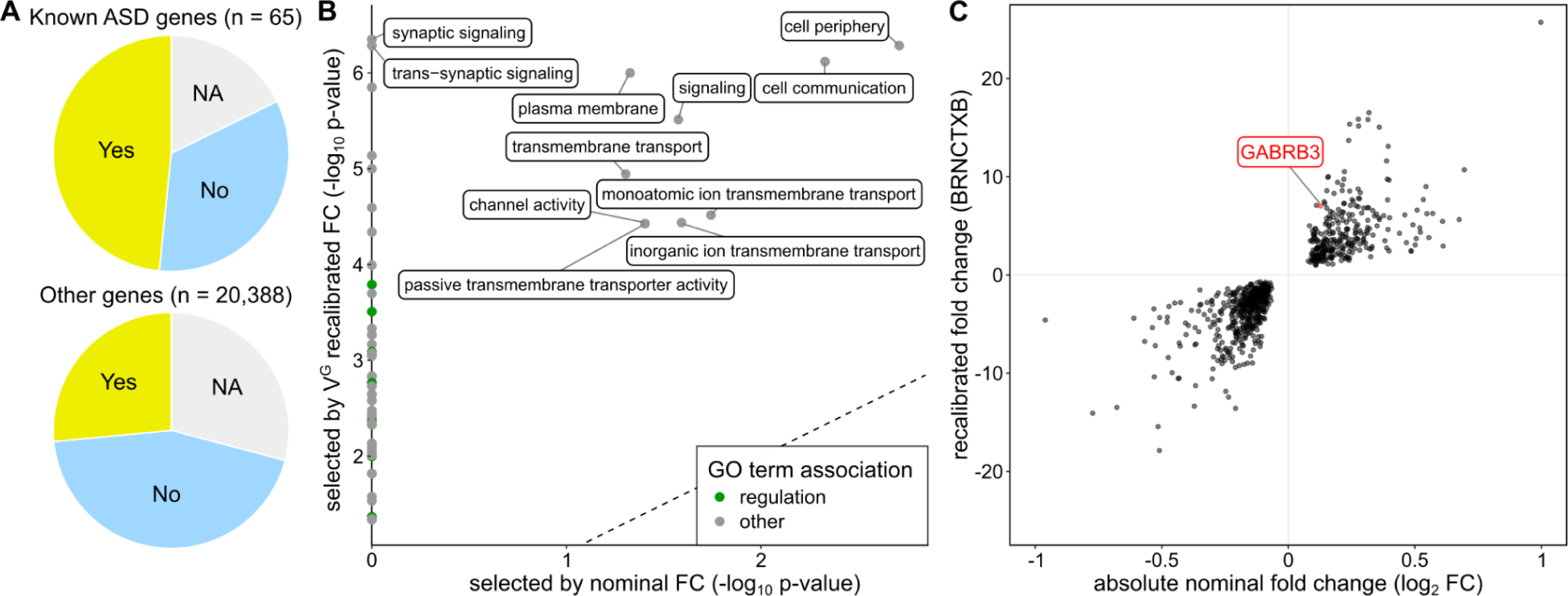
Tissue-specific recalibration in ASD. **A)** Comparison in which tissue the V^G^ of a gene is lowest. Compared to all other genes, known ASD genes are significantly more likely to have the lowest V^G^ in one of the GTEx brain tissues. NA are genes were no V^G^ value exists for any of the 14 GTEx brain tissues. **B)** GO term enrichment of DE experiment comparing ASD patients and controls. The GO terms ‘synaptic signaling’ and ‘trans synaptic signaling’ are significantly enriched only post recalibration **C)** Absolute nominal compared to recalibrated fold changes per gene in the ASD experiment, with the ASD-associated gene GABRB3 highlighted.

The complete psychENCODE dataset contains 25,772 genes. For recalibration, we used the V^G^ values of the GTEx Brain-Frontal Cortex tissue. Of the 1,611 statistically significant DE genes in ASD versus controls, 746 are recalibrated. We observed a general increase in GO term enrichments among the top 300 genes due to recalibration, with patterns analogous to those in the IFNγ stimulus data (Fig. 4B). The largest effect is seen in the highly overlapping GO terms ‘synaptic signaling’ (GO:0099536) & ‘trans synaptic signaling’ (GO:0099537), which were not significantly enriched when selecting genes based on nominal fold changes. Synaptic signaling is well-characterized as a key process in ASD (Jiang et al. 2022). Notably, enrichments were generally higher when recalibration is done with brain V^G^ compared to the cross-tissue mean V^G^ (Supplemental Fig. S11, Supplemental Table S4). We observed similar increased enrichments of development related GO terms for SCZ (Supplemental Fig. S12) but found no significant GO term enrichment for BD after selecting genes with V^G^ estimates from the already small number of DE genes. Only three of the 65 genetically identified ASD genes (An et al. 2018) are statistically significantly expressed in the psychENCODE DE data. One of these three genes is GABRB3. GABRB3 is one of the genes classified under the GO term synaptic signaling, and has been extensively studied for its role in neurological disorders, including ASD. Of note, ASD has been strongly linked to a duplication of GABRB3 (Cook et al. 1998), while several studies have found no (Mahdavi et al. 2018; Noroozi et al. 2018) or only limited (Adak et al. 2021) evidence of association with frequent single nucleotide polymorphisms in the gene. This indicates that changes in dosage rather than structure are causing the association of GABRB3 with ASD. In the psychEncode ASD experiment, GABRB3 expression was increased with a log fold change of 0.12, ranking as the 858th most differentially expressed gene in terms of absolute nominal fold change (Fig. 4C). After recalibration, the gene is one of the 100 most DE genes (rank 85), illustrating how recalibration can highlight a known causal disease gene.

## DISCUSSION

Here, we have introduced a novel approach for one of the most common analyses in computational biology: ranking of differentially expressed genes. Reprioritization of DE results with a genetic variance metric has the advantage of providing a more robust and generalizable estimate of biological constraint on expression, that is not confounded by intermingled technical and biological noise in specific expression studies. Furthermore, using *genetic* variance to better interpret environmental or other *non-genetic* differences brings together two typically distinct areas of biological enquiry. While a simpler method (Starr et al. 2023) has been proposed for assessing whether an observed expression change is different from the natural variability, V^G^ is based on allelic fold changes (Mohammadi et al. 2017) that are measured in the same unit as DE fold change. Thus, this the first method to compare different genes from an experiment on the same, biologically interpretable scale. Furthermore, we have shown that tissue-specific recalibration further increases the enrichment of likely regulatory pathways involved and is a unique advantage over most other constraint-based approaches for gene prioritization.

We have shown that recalibration deprioritizes highly variable genes and removes the bias of DE results easily highlighting non-essential genes with high population variation. Reranking of DE genes led to increased enrichment of molecular processes related to regulation, and correctly identified key driver processes in neuropsychiatric traits. This indicates that our approach has the potential to identify phenotypic drivers, guiding downstream analyses towards genes and processes that are biologically causal. These would represent potential targets for interventions. Conversely, the standard DE approach is sensitive in picking up genes that reflect the response to the signal of interest. While such genes do not necessarily drive downstream processes, they may be informative, for example, as biomarkers of external stimulus. Thus, the standard and recalibrated DE analyses highlight different types of biology, both of which may be of interest. However, the ability to focus the downstream analysis to phenotypic drivers and deprioritize highly variable genes that merely reflect the response will be a valuable in many applications.

However, the approach presented here has some limitations. One of them is missing V^G^ estimates for many genes, especially in tissue-specific analyses. Our previous and new findings show that missing genes are generally of very low expression, which is the reason why allele-specific data/eQTLs could not be determined, and that low expressed genes are generally more variable and thus negatively affected by recalibration. Thus, missing genes are likely to be negatively correlated to functional importance, unlike for some genetics-based scores (Lek et al. 2016). Generating V^G^ for additional genes via machine learning clearly reduces this limitation. Furthermore, growing eQTL and allelic expression datasets will further improve the number of data for which V^G^ can be generated, particularly in specific tissues and cell types. While larger datasets will also improve the quality of V^G^ estimates, our results demonstrate that the current V^G^ scores are of sufficient quality for our straightforward recalibration to yield biologically meaningful signals. This evidence will hopefully inspire future incorporation of V^G^ noise estimates in the fold change recalibration step, and potentially even into DE significance testing itself to account for general population variance. In such analyses, accurate matching of the cell type and sometimes even the population background between the experiment and V^G^ values is likely to become an even higher priority.

In summary, we have introduced a new approach for prioritizing potential regulatory drivers in differential expression analyses. This provides yet another demonstration of the value of population-scale RNA-sequencing data for enhancing biological interpretability of broadly used study designs and enabling new discoveries in the future.

## METHODS

### Genetic variance in gene expression

V^G^ is a metric that estimates the genetic variance in expression for genes in the human population, which we introduced in previous publications (Mohammadi et al. 2017, 2019). There, we described two approaches based on different data types to calculate V^G^: from allelic expression (AE) and from expression quantitative trait loci (eQTL). Briefly, AE-based V^G^ (V^G^_AE_) is derived based on the comparison of allele read counts for each single nucleotide variant (SNV) in the two haplotype copies in a diploid individual. This information is aggregated by SNV frequency over all individuals in the tested population and fitted via ANEVA (analysis of expression variation), a model for population AE data, to infer with V^G^_AE_ the level of genetic expression variability between haplotypes. In contrast, V^G^ is generated from the most significant eQTL of a gene as aggregate of effect size and allele frequency. Both approaches of generating V^G^ have been applied to RNA-Seq samples from the GTEx project, generating separate estimates for each GTEx tissue. A mean V^G^ has been calculated as a weighted harmonic mean over all tissue specific V^G^ estimates, using expression per tissue, measured in TPM, as weight.

V^G^ estimates used in this study were generated on data from GTEx v8 and were reused from the respective release by the GTEx Consortium (GTEx Consortium 2020). V^G^ estimates are based on GTEx v7 and were reused from the *Mohammadi et al. 2019* publication. Any V^G^ estimates used that are not further specified are AE-derived mean V^G^. Genes for which no mean V^G^ has been estimated were excluded from the non-machine learning analyses. All V^G^ estimates are provided in the supplement (Supplemental Table S5).

### Other gene metrics

RVIS scores per gene were obtained from Petrovski et al. 2013. ncCADD, ncGERP, and ncRVIS were obtained from *Petrovski et al. 2015*. The numbers of enhancers and super-enhancers per gene were derived from GeneHancer v5 (Fishilevich et al. 2017). Based on the gff file with all enhancers from genecards.org, enhancers with a score >= 0.7 and a gene association ‘link_score’ >= 7 were filtered. The final numbers represent the total counts after filtering per gene. The index for tissue specific expression in GTEx tau (Palmer et al. 2021), was downloaded from genomics.senescence.info/gene_expression/Tau_score.zip. The triplosensitivity and haploinsufficiency scores, pTriplo and pHaplo respectively, were obtained from the supplemental data of *Collins et al. 2022*. The same dataset was further used to obtain EpiScore (Han et al. 2018). pLI (Lek et al. 2016) and LOEUF (Karczewski et al. 2020) scores were obtained from gnomad.broadinstitute.org, using data from the v3 release. Transcripts per million (TPM) values for each tissue were downloaded from gtexportal.org, using GTEx version 8. When referring to TPM in the text, we are generally using the mean TPM across all GTEx tissues, which were in turn calculated as the median across all individual samples of each tissue.

To account for the different scales of different gene metrics, correlations with V^G^ were calculated as Spearman correlation. 95% confidence intervals of correlation were determined using bootstrapping.

The list of nonessential genes was obtained from *Zarrei et al. 2015*. ClinGen genes were downloaded via the dosage sensitivity curated gene list from ftp.clinicalgenome.org/ClinGen_gene_curation_list_GRCh38.tsv (version from 2023/02/13). Haploinsufficient genes were selected by joining all genes with a haploinsufficiency score of 1, 2 or 3. The background gene set contains all genes with ASE-based V^G^ values, except for the genes included in one of the two other gene sets.

### KEGG gene annotations

The annotation of KEGG functional categories was adapted from *Dong et al. 2023*. We reused their functional category labels, but excluded the category ‘Domain-containing proteins not elsewhere classified’ due to frequent overlaps with other categories. When multiple labels could be applied to a gene, we prioritized the one that appeared least frequently in the list of all genes with V^G^. Genes that were not found in KEGG or did not match any of the labels were omitted.

### Recalibrating expression fold change

In order to prioritize biological significance, expression fold change (FC) from differential expression analysis is standardized relative to the standard deviation of genetically regulated gene expression: 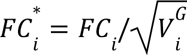, where *FC_i_* and *FC_i_*\* are the nominal and recalibrated log fold change of a gene *i*, and *V*^*G*^*_i_* is the expected variance introduced in gene dosage by genetic variation in the population.

### RNA expression data processing

RNA expression data from *Alasoo et al. 2018* was downloaded from zenodo.org/record/839011 (Alasoo 2017) and zenodo.org/record/4678936 (Alasoo and Kerimov 2021). The datasets contain the results after differential gene expression testing with DESeq2 and count data respectively. Differential expression was determined using an adjusted p-value of smaller 0.001 as significance cut-off. Recalibration was performed using the mean V^G^_AE_ values.

The *Findley et al. 2021* dataset of different perturbation agents were downloaded from the supplemental data of the publication. The experiments mentioned in the analysis are ‘15C1’ (copper perturbation) and ‘20C1’ (zinc perturbation) of iPSCs from the shallow sequencing dataset. The downloaded datasets contain the results of DE testing with DESeq2. Differential expression was determined using a false discovery rate cut-off of 0.05. Recalibration was performed using the mean V^G^_AE_ values.

The psychENCODE datasets from *Gandal et al. 2018* for ASD, SCZ and BD were downloaded from the supplementary data. The datasets contain the results after DE testing with DESeq2. Differential expression was determined using a false discovery rate cut-off of 0.05. Recalibration was performed using the machine learning assisted V^G^_AE_ values for BRNCTXB.

### Correlation between nominal and recalibrated fold changes

In order to avoid inflation caused by directionality (which does not change), we calculated the correlation between nominal and recalibrated fold changes using the Pearson method of the absolute values.

### GO term enrichment

Gene Ontology (GO) term enrichment analysis was conducted using the R package gprofiler2 (Raudvere et al. 2019; Kolberg et al. 2023), using GO terms and gene associations from the Biological Process (BP), Cellular Component (CC) and Molecular Function (MF) GO resources based on Ensembl release 110. Enrichments were calculated based on Ensembl gene identifiers against all genes with determined expression and V^G^ values in the dataset. Genes were selected by absolute nominal fold change and absolute recalibrated fold change respectively. For all enrichment comparisons except the testing of other gene numbers, the top ∼50% (rounded to the nearest 100) of all significant genes are selected. Unless otherwise stated, only genes from the experiment for which V^G^ has been calculated were included as the background set. Enrichment was considered up to an adjusted p-value of 0.05. In the comparison of nominal and recalibrated fold changes, non-enriched GO terms were imputed with a p-value of 1.

GO term clustering was performed with the help of the simplifyEnrichment package (Gu and Hübschmann 2023) using binary-cut clustering. Clustering had to be performed separately for terms of each GO resource (BP, CC & MF).

### Gene Set Enrichment

Gene Set Enrichment Analysis (GSEA) was performed using the R package clusterProfiler (Wu et al. 2021) using fgsea (Korotkevich et al. 2021) as backend. In contrast to GO term enrichment based on groups, only GO terms and gene association from the GO Biological Process resource were used, and the analysis was performed on all genes (irrespective of statistical significance) for which gene expression and V^G^ are defined. Enrichment was considered up to an adjusted p-value of 0.05. Corresponding non-enriched GO terms were imputed with a p-value of 1.

### Training a supervised machine learning model of V^G^

All data preparation for machine learning was done in Python, using scikit-learn. 14 features (Supplemental Table S2) were collected to train a supervised machine learning model of V^G^_AE_. All 14 features were available for 12,235 genes, of which 9,562 also had V^G^_AE_. The estimates of V^G^_AE_ and V^G^_AE_ were transformed by natural logarithm (ln). To increase the number of genes in our dataset, missing gene features were imputed using a K-nearest neighbor imputer (Troyanskaya et al. 2001), imputing genes with up to 5 missing features. 16,943 genes with all 14 features were obtained, of which 12,004 had estimates of V^G^_AE_. For model training, these were split in an 80% train and 20% hold-out test set. Gradient boosting trees were trained using the Python bindings of XGBoost (Chen and Guestrin 2016). Model parameters were optimized with hyperopt (Bergstra et al. 2013) and selected by minimal root mean squared error on 10-fold cross-validation. The final model parameters were max_depth = 6, learning_rate = 0.0236, subsample = 0.96, colsample_bytree = 0.746, colsample_bylevel = 0.494, n_estimators=344, min_split_loss = 2, reg_lambda = 1.27 and reg_alpha = 4.5. V^G^ values were generated for all 16,943 genes with the imputed feature set.

Tissue-specific models were trained separately for each tissue using the same training parameters as for V^G^_ML_. Tissue-specific V^G^_AE_ were used as labels, while TPM and V^G^_eQTL_ were used as tissue-specific features. To increase the number of predicted genes, existing tissue V^G^_AE_ were combined with the generated ones, with existing scores taking precedence if both existed for a particular gene. All V^G^_AEML_ are available in Supplemental Table S6.

The comparison of the number of genes per type of V^G^ (Upset plot) was generated with UpSetR (Conway et al. 2017).

### ASD genes

Previously ASD associated genes were downloaded from *An et al. 2018*, using the most stringent list with FDR<0.1. The tissue with the lowest V^G^ per gene was calculated from V^G^_AEML_, considering only tissues with a calculated V^G^ score for each gene. GTEx brain tissues are all 13 GTEx tissues starting with ‘BRN’ plus pituitary. Enrichment was determined using a one-sided binomial test.

## DATA ACCESS

All V^G^_AE_ estimates are provided in Supplemental Table S5. All V^G^_AEML_ estimates are provided in Supplemental Table S6. An R package for recalibrating experimental DE data is available at github.com/LappalainenLab/recalibrate.

## ACKNOWLEDGMENTS

We would like to thank Sarah Kim-Hellmuth, Xiaoting Li, Ryan Collins, Sanna Gudmundsson, Paul Hoffman, Mariia Mineava, Kaushik Ram Ganapathy, and the current and former members of the Lappalainen Lab for helpful discussions and code sharing. Part of the computations were enabled by resources provided by the Swedish National Infrastructure for Computing (SNIC) at UPPMAX partially funded by the Swedish Research Council through grant agreement no. 2018-05973.

## REFERENCES

Adak P, Sinha S, Banerjee N. 2021. An Association Study of Gamma-Aminobutyric Acid Type A Receptor Variants and Susceptibility to Autism Spectrum Disorders. J Autism Dev Disord 51: 4043–4053.

Alasoo K. 2017. Differential gene expression in iPSC-derived macrophages after IFNg stimulation and Salmonella infection. https://zenodo.org/record/839011 (Accessed October 3, 2023).

Alasoo K, Kerimov N. 2021. Public RNA-seq count matrices and sample metadata from the eQTL Catalogue. https://zenodo.org/record/4678936 (Accessed October 9, 2023).

Alasoo K, Rodrigues J, Mukhopadhyay S, Knights AJ, Mann AL, Kundu K, Hale C, Dougan G, Gaffney DJ. 2018. Shared genetic effects on chromatin and gene expression indicate a role for enhancer priming in immune response. Nat Genet 50: 424–431.

Albert FW, Kruglyak L. 2015. The role of regulatory variation in complex traits and disease. Nat Rev Genet 16: 197–212.

An J-Y, Lin K, Zhu L, Werling DM, Dong S, Brand H, Wang HZ, Zhao X, Schwartz GB, Collins RL, et al. 2018. Genome-wide de novo risk score implicates promoter variation in autism spectrum disorder. Science 362: eaat6576.

Benjamini Y, Hochberg Y. 1995. Controlling the false discovery rate: a practical and powerful approach to multiple testing. J R Stat Soc Ser B Methodol 57: 289–300.

Bergstra J, Yamins D, Cox D. 2013. Making a Science of Model Search: Hyperparameter Optimization in Hundreds of Dimensions for Vision Architectures. In Proceedings of the 30th International Conference on Machine Learning, pp. 115–123, PMLR https://proceedings.mlr.press/v28/bergstra13.html (Accessed October 20, 2023).

Chen T, Guestrin C. 2016. XGBoost: A Scalable Tree Boosting System. In Proceedings of the 22nd ACM SIGKDD International Conference on Knowledge Discovery and Data Mining, pp. 785–794 http://arxiv.org/abs/1603.02754 (Accessed July 27, 2023).

Collins RL, Glessner JT, Porcu E, Lepamets M, Brandon R, Lauricella C, Han L, Morley T, Niestroj L-M, Ulirsch J, et al. 2022. A cross-disorder dosage sensitivity map of the human genome. Cell 185: 3041–3055.e25.

Conway JR, Lex A, Gehlenborg N. 2017. UpSetR: an R package for the visualization of intersecting sets and their properties ed. J. Hancock. Bioinformatics 33: 2938–2940.

Cook EH, Courchesne RY, Cox NJ, Lord C, Gonen D, Guter SJ, Lincoln A, Nix K, Haas R, Leventhal BL, et al. 1998. Linkage-disequilibrium mapping of autistic disorder, with 15q11-13 markers. Am J Hum Genet 62: 1077–1083.

Cui X, Churchill GA. 2003. Statistical tests for differential expression in cDNA microarray experiments. Genome Biol 4: 210.

Dong D, Shen H, Wang Z, Liu J, Li Z, Li X. 2023. An RNA-informed dosage sensitivity map reflects the intrinsic functional nature of genes. Am J Hum Genet 110: 1509–1521.

Findley AS, Monziani A, Richards AL, Rhodes K, Ward MC, Kalita CA, Alazizi A, Pazokitoroudi A, Sankararaman S, Wen X, et al. 2021. Functional dynamic genetic effects on gene regulation are specific to particular cell types and environmental conditions. eLife 10: e67077.

Fishilevich S, Nudel R, Rappaport N, Hadar R, Plaschkes I, Iny Stein T, Rosen N, Kohn A, Twik M, Safran M, et al. 2017. GeneHancer: genome-wide integration of enhancers and target genes in GeneCards. Database 2017. https://academic.oup.com/database/article/doi/10.1093/database/bax028/3737828 (Accessed October 17, 2023).

Flynn ED, Lappalainen T. 2022. Functional Characterization of Genetic Variant Effects on Expression. Annu Rev Biomed Data Sci 5: 119–139.

Gandal MJ, Zhang P, Hadjimichael E, Walker RL, Chen C, Liu S, Won H, van Bakel H, Varghese M, Wang Y, et al. 2018. Transcriptome-wide isoform-level dysregulation in ASD, schizophrenia, and bipolar disorder. Science 362: eaat8127.

Ge Y, Sealfon SC, Speed TP. 2009. Multiple testing and its applications to microarrays. Stat Methods Med Res 18: 543–563.

Greenland S, Senn SJ, Rothman KJ, Carlin JB, Poole C, Goodman SN, Altman DG. 2016. Statistical tests, P values, confidence intervals, and power: a guide to misinterpretations. Eur J Epidemiol 31: 337–350.

GTEx Consortium. 2020. The GTEx Consortium atlas of genetic regulatory effects across human tissues. Science 369: 1318–1330.

Gu Z, Hübschmann D. 2023. simplifyEnrichment: A Bioconductor Package for Clustering and Visualizing Functional Enrichment Results. Genomics Proteomics Bioinformatics 21: 190–202.

Han X, Chen S, Flynn E, Wu S, Wintner D, Shen Y. 2018. Distinct epigenomic patterns are associated with haploinsufficiency and predict risk genes of developmental disorders. Nat Commun 9: 2138.

Harrison PF, Pattison AD, Powell DR, Beilharz TH. 2019. Topconfects: a package for confident effect sizes in differential expression analysis provides a more biologically useful ranked gene list. Genome Biol 20: 67.

Ignatiadis N, Klaus B, Zaugg JB, Huber W. 2016. Data-driven hypothesis weighting increases detection power in genome-scale multiple testing. Nat Methods 13: 577–580.

Jiang C-C, Lin L-S, Long S, Ke X-Y, Fukunaga K, Lu Y-M, Han F. 2022. Signalling pathways in autism spectrum disorder: mechanisms and therapeutic implications. Signal Transduct Target Ther 7: 229.

Jung K, Friede T, Beißbarth T. 2011. Reporting FDR analogous confidence intervals for the log fold change of differentially expressed genes. BMC Bioinformatics 12: 288.

Karczewski KJ, Francioli LC, Tiao G, Cummings BB, Alföldi J, Wang Q, Collins RL, Laricchia KM, Ganna A, Birnbaum DP, et al. 2020. The mutational constraint spectrum quantified from variation in 141,456 humans. Nature 581: 434–443.

Kolberg L, Raudvere U, Kuzmin I, Adler P, Vilo J, Peterson H. 2023. g:Profiler—interoperable web service for functional enrichment analysis and gene identifier mapping (2023 update). Nucleic Acids Res 51: W207–W212.

Korotkevich G, Sukhov V, Budin N, Shpak B, Artyomov MN, Sergushichev A. 2021. Fast gene set enrichment analysis. 060012. https://www.biorxiv.org/content/10.1101/060012v3 (Accessed December 2, 2022).

Lek M, Karczewski KJ, Minikel EV, Samocha KE, Banks E, Fennell T, O’Donnell-Luria AH, Ware JS, Hill AJ, Cummings BB, et al. 2016. Analysis of protein-coding genetic variation in 60,706 humans. Nature 536: 285–291.

Love MI, Huber W, Anders S. 2014. Moderated estimation of fold change and dispersion for RNA-seq data with DESeq2. Genome Biol 15: 550.

Mahdavi M, Kheirollahi M, Riahi R, Khorvash F, Khorrami M, Mirsafaie M. 2018. Meta-Analysis of the Association between GABA Receptor Polymorphisms and Autism Spectrum Disorder (ASD). J Mol Neurosci 65: 1–9.

McCarthy DJ, Smyth GK. 2009. Testing significance relative to a fold-change threshold is a TREAT. Bioinformatics 25: 765–771.

Mohammadi P, Castel SE, Brown AA, Lappalainen T. 2017. Quantifying the regulatory effect size of cis-acting genetic variation using allelic fold change. Genome Res 27: 1872–1884.

Mohammadi P, Castel SE, Cummings BB, Einson J, Sousa C, Hoffman P, Donkervoort S, Jiang Z, Mohassel P, Foley AR, et al. 2019. Genetic regulatory variation in populations informs transcriptome analysis in rare disease. Science 366: 351–356.

Mostafavi H, Spence JP, Naqvi S, Pritchard JK. 2023. Systematic differences in discovery of genetic effects on gene expression and complex traits. Nat Genet 55: 1866–1875.

Naqvi S, Kim S, Hoskens H, Matthews HS, Spritz RA, Klein OD, Hallgrímsson B, Swigut T, Claes P, Pritchard JK, et al. 2023. Precise modulation of transcription factor levels identifies features underlying dosage sensitivity. Nat Genet 55: 841–851.

Noroozi R, Taheri M, Ghafouri-Fard S, Bidel Z, Omrani MD, Moghaddam AS, Sarabi P, Jarahi AM. 2018. Meta-analysis of GABRB3 Gene Polymorphisms and Susceptibility to Autism Spectrum Disorder. J Mol Neurosci 65: 432–437.

Palmer D, Fabris F, Doherty A, Freitas AA, de Magalhães JP. 2021. Ageing transcriptome meta-analysis reveals similarities and differences between key mammalian tissues. Aging 13: 3313–3341.

Petrovski S, Gussow AB, Wang Q, Halvorsen M, Han Y, Weir WH, Allen AS, Goldstein DB. 2015. The Intolerance of Regulatory Sequence to Genetic Variation Predicts Gene Dosage Sensitivity. PLOS Genet 11: e1005492.

Petrovski S, Wang Q, Heinzen EL, Allen AS, Goldstein DB. 2013. Genic Intolerance to Functional Variation and the Interpretation of Personal Genomes. PLOS Genet 9: e1003709.

Raudvere U, Kolberg L, Kuzmin I, Arak T, Adler P, Peterson H, Vilo J. 2019. g:Profiler: a web server for functional enrichment analysis and conversions of gene lists (2019 update). Nucleic Acids Res 47: W191–W198.

Rice AM, McLysaght A. 2017. Dosage-sensitive genes in evolution and disease. BMC Biol 15: 78.

Ritchie ME, Phipson B, Wu D, Hu Y, Law CW, Shi W, Smyth GK. 2015. limma powers differential expression analyses for RNA-sequencing and microarray studies. Nucleic Acids Res 43: e47.

Robinson MD, McCarthy DJ, Smyth GK. 2010. edgeR: a Bioconductor package for differential expression analysis of digital gene expression data. Bioinformatics 26: 139–140.

Schena M, Shalon D, Davis RW, Brown PO. 1995. Quantitative monitoring of gene expression patterns with a complementary DNA microarray. Science 270: 467–470.

Sekovanić A, Jurasović J, Piasek M. 2020. Metallothionein 2A Gene Polymorphisms in Relation to Diseases and Trace Element Levels in Humans. Arch Ind Hyg Toxicol 71: 27–47.

Starr AL, Gokhman D, Fraser HB. 2023. Accounting for cis-regulatory constraint prioritizes genes likely to affect species-specific traits. Genome Biol 24: 11.

Troyanskaya O, Cantor M, Sherlock G, Brown P, Hastie T, Tibshirani R, Botstein D, Altman RB. 2001. Missing value estimation methods for DNA microarrays. Bioinformatics 17: 520–525.

Wasserstein RL, Lazar NA. 2016. The ASA Statement on p-Values: Context, Process, and Purpose. Am Stat 70: 129–133.

Wu T, Hu E, Xu S, Chen M, Guo P, Dai Z, Feng T, Zhou L, Tang W, Zhan L, et al. 2021. clusterProfiler 4.0: A universal enrichment tool for interpreting omics data. The Innovation 2: 100141.

Zarrei M, MacDonald JR, Merico D, Scherer SW. 2015. A copy number variation map of the human genome. Nat Rev Genet 16: 172–183.

Zhang S, Cao J. 2009. A close examination of double filtering with fold change and t test in microarray analysis. BMC Bioinformatics 10: 402.

